# Unbiased analysis of *C. elegans* behavior reveals the use of distinct turning strategies during magnetic Orientation

**DOI:** 10.1101/688408

**Authors:** C Bainbridge, J McDonald, A. Ahlert, Z Benefield, W Stein, AG Vidal-Gadea

## Abstract

To successfully navigate their surroundings, animals detect and orient to environmental stimuli possessing unique physical properties. Most animals can derive directional information from spatial or temporal changes in stimulus intensity (e.g. chemo- and thermo-taxis). However, some biologically relevant stimuli have constant intensity at most organismal scales. The gravitational and magnetic fields of the earth are examples of uniform stimuli that remain constant at most relevant scales. While devoid of information associated with intensity changes, the vectorial nature of these fields intrinsically encodes directional information. While much is known about behavioral strategies that exploit changes in stimulus intensity (gradients), less is understood about orientation to uniform stimuli. Nowhere is this truer than with magnetic orientation. While many organisms are known to orient to the magnetic field of the earth, how these animals extract information from the earth’s magnetic field remains unresolved.

Here we use the nematode *C. elegans* to investigate behavioral strategies for orientation to magnetic fields, and compare our findings to the better characterized chemical and thermal orientation strategies. We used an unbiased cluster analysis to categorize, quantify, and compare behavioral components underlying different orientation strategies as a way to quantify and compare animal orientation to distinct stimuli. We find that in the presence of an earth-like magnetic field, worms perform acute angle turns (140-171°) that significantly improved their alignment with the direction of an imposed magnetic vector. In contrast, animals performed high amplitude turns (46-82°) that significantly increased alignment of their trajectory with the preferred migratory angle. We conclude that *C. elegans* orients to earth-strength magnetic fields using two independent behavioral strategies, in contrast to orientation strategies to graded stimuli. Understanding how *C. elegans* detects and orients to magnetic fields will provide useful insight into how many species across taxa accomplish this fascinating sensory feat.

## Background

Animals sense and integrate different environmental stimuli in order to optimize conditions for growth, survival, and reproduction. Locomotion in stimulus-rich habitats is particularly challenging since distinct physical parameters need to be simultaneously sensed, processed, and evaluated. Animals must therefore make use of different stimulus properties, extracting directional information, and selecting appropriate behavioral strategies in order to successfully navigate their environment.

For many natural stimuli, directional information is encoded in spatial or temporal changes intensity (gradients) that can vary linearly or radially away from a source (e.g. chemical gradients, Fig 1A). Behavioral strategies that allow animals to effectively orient to graded stimuli of different natures have been thoroughly studied (e.g. visual: el Jundi et al., 2014; Ruppertsberg et al., 2008; auditory: Teder-Sälejärvi and Hillyard, 1998; chemical: Pierce-Shimomura et al., 1999; and thermal: Kimata et al. 2012). While behavioral strategies may differ depending on the type and shape of graded stimuli, they are similar in that information is derived from experienced spatiotemporal changes in stimulus intensity. Such strategies are usually limited to the narrow effective ranges over which animals are able to detect and compare intensity changes. While variable at planetary and geological scales, at organismal scales the magnetic field of the earth is largely devoid of temporal or spatial variation. Because of its vectorial nature, the earth’s field intrinsically possesses directional information. For animals able to detect it, this force field offers constant and reliable navigational information that must be detected without relying on spatial or temporal intensity changes.

**Figure 1.**
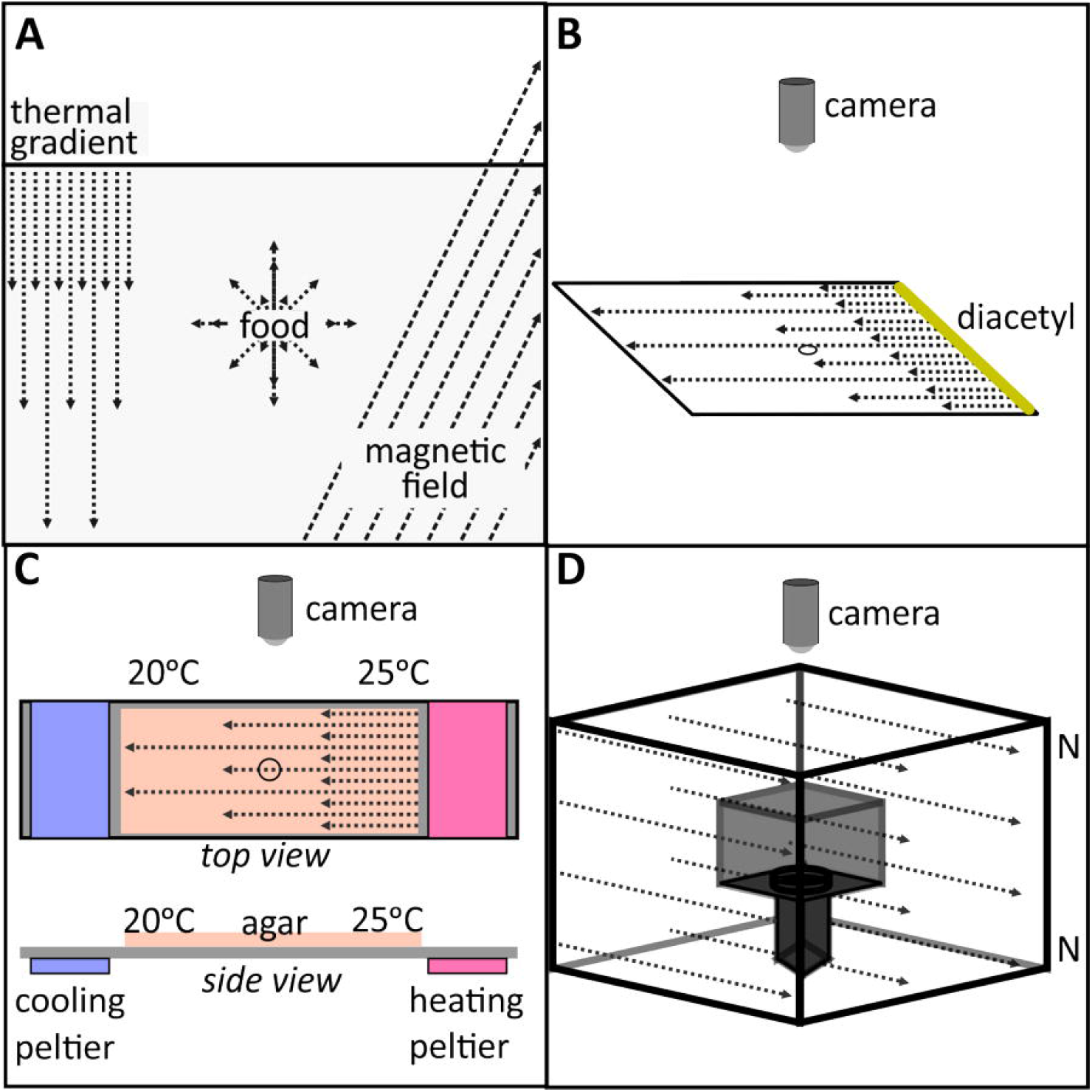
Experimental setup. **(A)** *C. elegans* orients to diverse stimuli in its environment which range from linear and radial gradients to uniform fields. **(B)** To assess chemotaxis an attractant (Diacetyl, yellow solid line) was applied along one edge of assay. Worms were placed at the start position (open circle) and filmed for 30 minutes. **(C)** To assess thermotaxis animals were cultivated at 25°C. A 5°C/cm thermal gradient was established using a pair of TECs to heat or cool the agar. **(D)** A double-wrapped 3D magnetic Merritt coil system composed of independently powered was used to generate uniform magnetic fields.

Animals are known to navigate using different vectorial parameters associated with the earth’s magnetic field (e.g. inclination and polarity, Wiltschko and Wiltschko, 2005). For example, European robins and sea turtles use magnetic field inclination to aid navigation (Wiltschko and Wiltschko, 1972, Lohmann et al., 2012; Lohmann and Lohmann, 1994; Schwarze et al., 2016). Alternatively, some fish species rely instead on the field’s polarity during migrations (Putman et al., 2014). While studies like these studies made great strides identifying how animals use magnetic fields, much work remains to be done to understand the behavioral strategies underlying orientation. Unlike transient stimuli, the magnetic field of the earth is always present. This necessitates that any adaptive interaction between an animal and the magnetic field must be initiated and terminated by internal organismal drives (or states). Orientation to the magnetic field is therefore state-dependent, and often initiated in response to other variables (e.g. reproductive season, Putman et al., 2014; Avens and Lohmann, 2004; Kullberg et al., 2003).

We recently showed that the nematode *C. elegans* detects and orients to magnetic fields (magnetotaxes, Vidal-Gadea et al., 2015). Worms been effectively used to study animal locomotion (Gjorgjieva et al., 2014; Zhen and Samuel, 2015). They perform several discreet, stereotypical reorientation maneuvers when exploring their environments (Croll, 1975). Much progress has been made in understanding how *C. elegans* detect and orient to chemical and thermal gradients (Goodman et al., 2005; Hart and Chao, 2010). During chemotaxis, worms decrease the frequency of reorientation events (called pirouettes) when moving towards a stimulus; effectively increasing the likelihood of approaching an attractant (Pierce-Shimomura et al., 1999). A similar strategy, known as a random walk, occurs during bacterial chemotaxis (Berg, 1993). Additionally, *C. elegans* perform a behavior called weathervaning where they gradually steer their crawling trajectories toward the source of a stimulus (Iino and Yoshida, 2009). Thermotaxis in worms relies on comparing surrounding temperature (T_S_) to a learned cultivation temperature (T_C_). Compared to chemotaxis, worms orienting to thermal gradients employ different behavioral strategies to arrive at a preferred thermal zone (Jurado et al., 2010). The strategy adopted by thermotaxing worms differs whether animals are moving up or down a thermal gradient (Luo et al., 2014). Similarly to chemotaxis, when T_S_>T_C_, *C. elegans* decrease reorientations when headed toward T_C_. However, when T_S_<T_C_ worms only bias the direction of turns toward increasing temperature. The net effect of both behavioral strategies is to orient the animal in the direction of T_C_.

*C. elegans* use the primary thermosensory AFD neurons to detect magnetic stimuli (Vidal-Gadea et al., 2015). It remains unclear if the common cellular basis of detection of thermal and magnetic stimuli is reflected in the orientation strategies adopted by worms. Because of the shared cellular basis for sensory detection, temperature and magnetic orientation offer a unique opportunity to disambiguate differences in orientation strategies that reflect underlying neural substrates, versus those that reflect differences between the physical nature of the stimuli. We hypothesize that *C. elegans* performs different behavioral strategies to orient to graded stimuli (chemical and thermal), compared to orientation to vectorial stimuli (magnetic fields).

Here we filmed *C. elegans* worms freely behaving in the presence of chemical, thermal, magnetic, and no-stimulus controls. Animal trajectories were digitized using machine vision. We constructed and used a custom behavioral algorithm to objectively quantify behavioral strategies used by worms under different experimental conditions. Our results indicate that *C. elegans* use different strategies to orient to chemical and thermal gradients compared to magnetic fields.

## Materials and Methods

### Animal culture

Wild-type *C. elegans* (N2 strain) were obtained from the Caenorhabditis Genetics Center. Animals were raised under standard conditions (Brenner, 1974) at 20°C on nematode growth media (NGM) agar plates and fed *E. coli* (OP50). Environmental temperature and humidity can alter worm behavior. Thus, we only performed assays on animals raised at 20°C and between 30-37% relative humidity for all but the temperature experiments. Animals to be assessed for thermal orientation were instead cultivated at 25°C. All animals tested were never-starved, bleach-synchronized (Porta-de-la-Riva et al., 2012), day 1-adult hermaphrodites from never infected or overcrowded (<100 adults/plate) culture plates.

### Animal transfer

Between 20 and 30 worms were rinsed using liquid NGM and transferred using a 2μl volume as previously described (Bainbridge *et al., 2016*). We used a small piece of KimWipe to soak up excess liquid. This reduced the time required by the agar in the plate to absorb the remaining liquid and release the animals to begin the assay (as worms are unable to breach the surface tension of a liquid droplet). The time between removal from culture plate and the start of the assay (when worms are free of the transfer liquid) is a crucial experimental variable which we kept to under 5 minutes. To minimize other variables, we limited the number of assays run per culture plate to two.

### Immobilization of animals

For the purpose of tallying animal choice under different assay conditions, we used sodium azide to immobilize them. For our linear assays, 10μl of (100 mM) sodium azide (NaN_3_) were painted along two opposing edges of the plate (5 cm from the center of the plate). We allowed the agar to absorb the azide for 5 minutes before animals were added to the center of the plate as described above. For radial assays we placed 1μl 1M NaN_3_ over the stimulus, and another equal volume equidistantly on the opposite side of the start position. As above, we allowed 5 minutes before transferring animals to the assay center.

### Linear assay plates

All linear assays were performed in 10 cm square petri dishes containing 20mL of 3% chemotaxis agar (Wicks et al., 2000). Plates cured at 20°C and 37% humidity overnight. Linear gradients and controls were done in square plates, while radial gradients and controls were done using circular plates.

### Linear chemotaxis

We used 20μl of 0.1% diacetyl (TCI America, Inc.) to paint an attractant line one edge of the plate (Fig. 1B). We allowed the agar plate to absorb the diacetyl for 25 minutes before the start of the assays. Chemotaxis controls were run in plates with no diacetyl.

### Linear thermotaxis

We generated linear thermal gradients using a grounded 2D aluminum thermal stage with two (independently powered) thermoelectric cooling devices (TECs) as previously described (Daniels and McKemy, 2010). A 20mL slab of 3% chemotaxis agar was placed on the thermal stage. We allowed 20 minutes to establish a 0.5°C*cm^−1^ thermal gradient across the surface of the agar. We set the highest temperature to 25°C along one edge of the agar (cultivation temp), with 20°C on the opposite end (Fig. 1C). Animals were placed in the center at 23°C. Gradient steepness was confirmed during the assay by sampling the surface of agar with digital thermometers (Vernier, Beaverton, OR). Temperature controls were run in plates with no measurable temperature gradient across the length of the assay.

### Linear magnetotaxis

We used a triple Merritt coil system (Merritt et al., 1983) to generate uniform, linear, magnetic fields over the volume of the assay plates as previously described (Vidal-Gadea et al., 2019). The system is double wrapped so that the same amount of current may be passed in parallel (which generates two equal magnetic amplitude fields that sum in strength) or in the antiparallel configuration (which generates two magnetic fields of equal magnitude that cancel each other out, Fig. 1D). A Faraday cage constructed from copper mesh was placed and grounded around the sample to prevent electric fields from influencing animal behavior. We used the system in three configurations: **1) *Earth conditions***. The coil system was run in parallel configuration and used to generate a homogeneous and linear magnetic field of earth strength (0.65 Gauss). **2) *No-field control***. The system generated a magnetic field of the same strength but opposite direction as of earth’s local magnetic field. This effectively cancelled out the magnetic field of the earth within the test volume. **3) Current control**. To control for extraneous parameters potentially associated with magnetic field induction we ran current controls. Current controls were identical to *Earth condition* experiments, except that the coils were run in their antiparallel configuration. Therefore, while the same amount of current powered the system in the two conditions, under *current control* conditions two magnetic fields of equal strength but opposite polarity were generated. This effectively cancelled the magnetic stimulus generated but not any electric or heat noise potentially present. A milligauss magnetometer was used to setup and sample magnetic fields within the test volume to within 0.1mGauss (AlphaLab Inc., Salt Lake City).

### Radial assay plates

All radial assays were performed in 10cm circular petri dishes containing 20mL of chemotaxis agar. Stimuli were placed 1.7cm away from the center of the plate (start position) and allowed to incubate for 25 minutes before the start of the assay.

### Radial chemotaxis

To establish radial attractant geometry, we placed a 2μl drop of a 0.1% solution of diacetyl in water. Diacetyl was placed 1.7cm from the center start position. Diacetyl was allowed to be absorbed and diffuse into the agar for 20 minutes to establish the radial gradient.

### Radial thermotaxis

We placed a 0.5cm diameter aluminum rod controlled by a heated water bath 1.7cm from the center start position. The rod contacted the agar through a round 0.5cm window cut into the bottom of the assay plate. We used a digital thermometer to ensure a peak temperature of 25°C at the surface of the agar above the center of the heating rod. Temperature was monitored using digital thermometer.

### Radial magnetotaxis

We used N42 3.5-cm diameter neodymium magnet (K&J Magnetics Inc., Plumsteadville) to produce radial magnetic gradients as previously described (Vidal-Gadea et al., 2015). Magnets were positioned north side up under the assay dish so that the magnet center was 1.7cm to one side, and the edge of the magnet passed directly under the animal starting position. All radial magnetotaxis experiments were carried out in a temperature and humidity controlled environment (at 20°C and 37% humidity). The chamber was electrically isolated with a grounded Faraday cage to eliminate the influence of electric fields on animal behavior.

### Orientation index

Orientation indexes were calculated using the final positions of animals paralyzed by NaN3 at the end of the assays. This index was calculated as previously described (Wicks et al., 2000). Briefly, the orientation index is a value between zero and one and it is defined as: OI=[(T-C)/(T+C)], where T is the number of animals immobilized by test stimulus, and C is the number of animals immobilized by the opposite (control) site. Therefore an index of 1 indicates all animals migrated in the same direction, while an index of 0 means that animals migrated randomly with respect to the stimulus. A minimum of 10 assays were performed for each linear stimulus geometry, and 5 assays for each radial stimulus geometry.

### Behavior acquisition

We recorded animal behavior over a 36mm field of view centered at the start point using a USB microscope (Plugable, Redmond, WA) controlled by Micro-Manager Open Source Microscopy Software at a resolution of 640×480 pixels. Animals were illuminated using two overhead LED light sources and recorded for 30 minutes at 1hz and sub-sampled at a rate of 0.2Hz for analysis. Assays were randomly rotated between recordings to avoid influences from external variables on animal orientation. All videos used in this study will be available in peer-reviewed version of this manuscript.

### Animal tracking

Animal centroids were obtained and tracked using Image-Pro Plus 7 Software (Media Cybernetics, Rockville) as previously described (Vidal-Gadea et al., 2011). Briefly, the software uses machine vision to detect and track worm centroids and returns their x and y coordinates over time, as well as trajectories, instantaneous angles, and velocities. Worms were only tracked once they travelled at least 0.5 cm from the center starting position and only recordings that had at least 5 animals participating in the assay for more than 100 seconds were used for behavioral analyses. All assays consisted of 20-30 animals.

### Animal velocity

We calculated instantaneous velocity as the distance travelled between consecutive centroid position divided by the sampling time. Instantaneous velocities were averaged for each trajectory and reported as a mean velocity for the assay.

### Animal directional headings

To determine animal headings, we used a custom script written in Spike2 to analyze changes in animal position over time. Our script used animal centroid coordinate data to determine mean population headings (imported from Image-Pro Plus, see above). Worm responses to diverse stimuli change over time based on their internal state (Gosh et al., 2016; Witham et al., 2016; Klein et al., 2017). To compare animals in similar states, we binned each animal track into 5% intervals for each trajectory in order to normalize headings for animals that completed the same proportion of their total trajectory. It is important to note that our analysis is not based on the trajectory of migrating animals across the entire assay plates. Rather, our analysis is restricted to the initial trajectories in the central 3.6 cm region of the plate where we were able to resolve animals in order for our automated tracking software to detect and track their centroids.

Headings were determined by calculating the angle between the direction of locomotion (for each 5% track interval) relative to the shortest vector to the stimulus. Headings were reported out of 360 degrees, with 0° and 360° corresponding to headings directly toward the stimulus, 180° for headings directly away from the stimulus. <u>*Mean bin headings were*</u> calculated as the averaged headings across animals for each bin interval within an assay. Mean bin headings were averaged to obtain the population heading for an assay (<u>*assay heading*</u>).

### Turn angle distribution and criteria

A wealth of research on *C. elegans* locomotion has identified several turn strategies favored by the animal (Zhao et al., 2003; Gray et al., 2005; Kim et al., 2011). In order to perform automated and unbiased behavioral analysis, we restricted our script to detect and quantify centroid trajectory changes (rather than changes in animal body posture). Therefore, to identify and compare turning strategies performed during different behaviors, we wrote a script that calculated trajectory changes for each animal centroid between every consecutive video frame. A turn angle was determined by taking the interior angle (θ_i_) formed between three consecutive centroid positions, and then calculating the deflection from a straight path. Turn angles were grouped across all animals in all assay conditions to obtain a distribution of total turn angles used by all animals in all assay and experimental conditions (N=81,006 angles). Importantly, while our analysis identified trajectory changes (centroid turn angles) associated with the performance of known behaviors in *C. elegans* (e.g. reversals and weathervanes), our centroid trajectory analysis does not specifically assign body postures to changes in animal position.

We used a Gaussian mixture model (GMM using the Mclust R software package to determine discrete groupings of frequently performed turn angles by animals under all test conditions. GMMs allow for an unbiased and accurate model-based approach to estimate the density of data by fitting multiple gaussian components that describe the total probability density estimate (PDE, Najar et al., 2017). We used the PDE to identify distinct turn angle categories using the components of each gaussian fit. The categories identified through this approach reflected posture-based turning descriptions described in the literature.

We used the range of turn angles produced by our experimental population to define different turning behaviors or strategies. The optimal number of components describing our dataset was confirmed by Bayesian information criterion (BIC, Schwarz, 1978). We therefore used descriptive statistics from individual gaussian categories as criteria to define discrete turning behaviors. We used the interquartile ranges (IQR=25^th^-75^th^ percentile) of turn angles that fell under each Gaussian category. This allowed us to quantify specific behavioral components as a discrete intervals of turn angles. Turn angles falling outside of the defined IQRs were excluded from analysis. Behavioral detection was therefore performed using a custom algorithm based on corresponding turn angle intervals. The algorithm is available online (Bainbridge et al., 2019).

### Behavioral detection algorithm and rate analysis

Behavioral detection was performed using a custom algorithm written in Spike2. The algorithm used the IQR of turn angles from gaussian categories to define behavioral events in animal trajectories. From this detection algorithm we obtained the temporal and spatial coordinates for behavioral events. Behavioral rates were calculated from detected events within an animal’s trajectory divided by the trajectory duration. Mean behavioral rates were then averaged across animals in an assay. All statistics are reported as the mean behavioral rate by assay (min^−1^± s.e.m.).

### Spatial distribution of behaviors

To determine if animals alter the performance of different behaviors based on their distance to the stimulus, we obtained a heatmap of total animal positions within each stimulus condition. Heat maps were derived from 2D probability density estimate (PDE) of spatial coordinates from animal trajectories using the R-software package. This allowed us to determine animal spatial distribution throughout each assay. We next obtained similar heat maps for each behavioral component. This allowed us to determine spatial distributions of separate behavioral components. Since the likelihood of detecting a behavioral component is greatest where animal trajectories are the most dense, we normalized the PDE of detected behaviors to the PDE of animal trajectories. Normalization was done by dividing the total number of behavioral events detected by the total number of animal positions in heatmap spatial bins. This allowed us to determine a new PDE of where behavior densities were reported independently from the effect of position density.

We confirmed our spatial distribution findings by horizontally binning the distance to stimulus into 10 horizontal bins. Each bin was used to determine the fraction of total trajectory positions that corresponded to individual behavioral components. This measure was referred to as the track fraction. For example, *acute angle turn* track fractions were calculated as the total number of acute angle turns in a horizontal bin divided by the total number of animal positions in that bin. We then determined if track fractions were significantly correlated with binned distance to stimulus. A significant correlation indicated a change in turn probability with distance from stimulus. We reported mean track fractions for horizontal bins for each assay. Significance was determined using Pearson product moment correlation. Data outside of the 99% prediction interval were excluded.

### Turning bias analysis

To determine if the performance of turning behaviors helped animals orient toward a stimulus, we performed an analysis of turning bias. Directional turning bias was determined by calculating animal headings immediately before and after performance of a behavioral component. Animal headings before a turn were calculated from an animal’s previous position to the position of the detected behavior. Headings after a behavior were determined from the position of the detected behavior to the animal’s next position. Headings before or after a behavior were calculated independently. We reported ***mean headings*** before and after each behavioral component for animals by assay.

### Sample Sizes and Statistical Analysis

Analysis was performed on recordings for control (N=10), chemical (N=10), thermal (N=13), magnetic (N=10), current control (N=13), and no field (N=11) assays.

Significance for behavioral rates were determined with one-way ANOVA or nonparametric Kruskal-Wallis ANOVA on ranks when normality or equal variance requirements were not met. Rates of behaviors were compared post-hoc to control rates of behaviors using Holm-Sidak or Dunn’s corrections for post-hoc comparisons of parametric or non-parametric data respectively. We report mean±s.e.m for parametric tests and median±s.e.m values from ranks for non-parametric tests. Significance of linear correlations for spatial distributions were determined using Pearson product moment correlation on mean track fractions and reported as the correlation coefficient and resulting p-value. Behavioral rate graphs and linear regressions were plotted using SigmaPlot (Systat) software. Heatmaps were generated using R statistical software (ggplot package).

### Circular Statistics

Circular statistical analysis for animal headings were performed in the circular statistics toolbox (CircStat2012a, Berens, 2009) in Matlab R2018a (Mathworks, Natick MA). Significance of animal headings and headings before and after behavioral events were determined using Rayleigh’s test for non-uniformity. Significant p-values indicate animal headings were in a particular direction that differed from a random Von-Mises circular distribution. Headings toward (or aligned with) a stimulus were defined to be 0°, while those directly away as having 180°. Radius vector length (r-value) of animal headings corresponded to clustering of animal headings in a particular direction. Thus, r-values reflected strength of population heading response. For example, if the entire population displayed the same migratory angle, their *r* value would be r=1 (1= radius), but if the population migration was randomly distributed, then r=0.

## Results

To determine if *C. elegans* alters its behavioral strategies to orient to categorically distinct sensory stimuli, we tracked wild-type animals as they oriented towards chemical, thermal, and magnetic stimuli (Fig. 1B-D, Sup Video). Because in nature the geometry of these stimuli differ from one another (e.g. linear magnetic fields vs radial chemical gradients) we used linear and radial stimulus geometries. Animals placed in radial gradients showed significant orientation indices (OI) for chemical (OI=0.86±0.05sem, p<0.05, Holm-Sidak), thermal (OI=0.89±0.07sem, p<0.05, Holm-Sidak), and magnetic stimuli (OI=0.39±0.06sem, p<0.05, Holm-Sidak) compared to controls (N=37, F(3,33)=32.98, one-way ANOVA p<0.001, Fig 2A). These findings were also observed when the stimuli were presented as linear gradients, where the one-way ANOVA showed significant difference between tests and controls (Fig. 2B, Pooled: N=51, F(3,48)=25.11, p<0.001). Unlike the random distribution of controls (OI=0.10±0.03sem), chemotaxing animals showed robust orientation toward the attractant (OI=0.87±0.05sem, p<0.05, Holm-Sidak post-hoc). Similarly, animals in a thermal gradient migrated towards their cultivating temperature (OI=0.55±0.05sem, p<0.05, Holm-Sidak post-hoc). Consistent with our previous work (Vidal-Gadea et al., 2015), animals migrating in a linear magnetic field of earth-strength oriented to the stimulus by aggregating at a preferred direction with respect to the applied field (OI=0.45±0.08sem, p<0.05, Holm-Sidak), showing a statistically significant migratory behavior (p<0.05, Dunn’s test) which was significantly different (Kruskal-Wallis ANOVA on ranks, H(2)=15.26, p<0.001) from animals in *no-field* (OI=0.14±0.02sem), and *current* (OI=0.13±0.03 sem) control conditions (Fig. S1 A). Because radial and linear conditions exhibited similar orientation indices, and because the geomagnetic field is uniformly linear at organismal scales, we elected to focus our analysis on stimuli of linear geometries.

**Figure 2.**
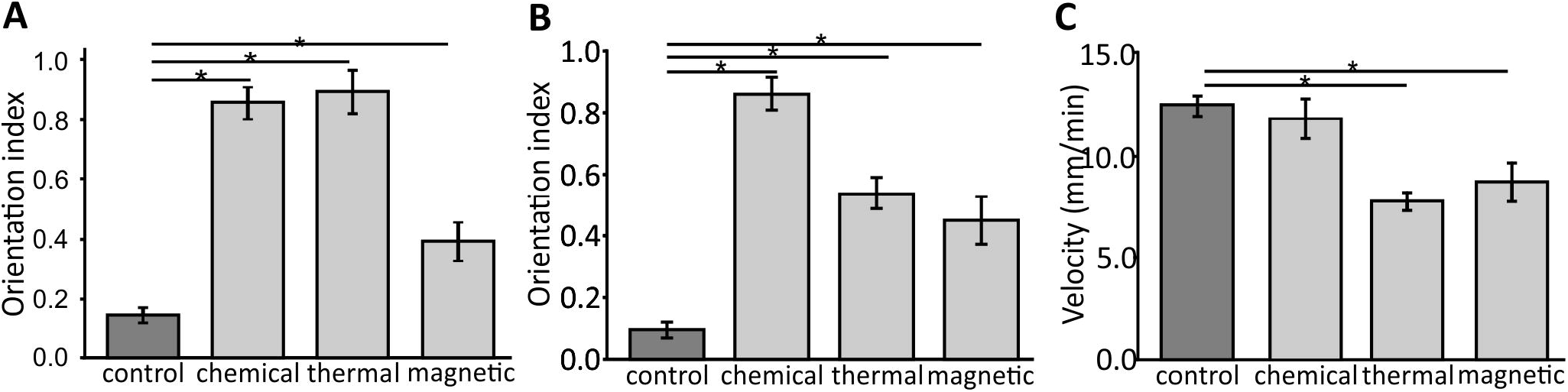
Animals orienting to thermal and magnetic stimuli decrease their crawling velocity. **(A)** Orientation indices for animals orienting. **(B)** Orientation indices for animals to linear chemical, thermal, and magnetic stimuli. Asterisks correspond to p<0.05 using Holm-Sidak post hoc correction. **(C)** Mean crawling velocity of animals migrating to linear chemical, thermal, and magnetic stimuli. Migrating animals orienting to thermal and magnetic stimuli reduced their mean velocity (asterisk indicates p<0.05).

### Animals reduced their crawling velocity when orienting to thermal and magnetic stimuli

Having established the three different orientation behaviors in our assays, we turned to analyze gross differences between the behaviors. Comparisons of animal velocities using centroid trajectories showed a significant velocity change between test and control animals (Fig. 2C, One-way Anova, N=42, F(3,39)=10.7, p<0.001). Compared to control animals (12.3mm*min^−1^±0.5sem), crawling velocities were significantly slower during thermotaxis and magnetotaxis (7.7mm*min^−1^±0.4sem and 8.6mm*min^−1^±0.9sem respectively; p<0.05, Holm-Sidak for each). However, animals orienting to chemical stimuli (chemotaxis) showed no significant crawling velocity difference from controls (11.7mm*min^−1^±1sem). These findings suggest that orientation to thermal and magnetic stimuli may place similar sensory demands on *C. elegans*, different from those imposed by chemical stimuli.

We next turned to analyze the population trajectories during the different experimental conditions (Fig. 3A). For this analysis we used circular statistics and described the population average heading relative to the stimulus (Fig. 3B, methods). Under control conditions, animals showed no significant directional heading (angle=306.0°, Rayleigh test: p=0.14, r=0.45). However, animals orienting to chemical and thermal gradients displayed a migration biased toward an attractant (angle=6.7°, Rayleigh test = p<0.01, r=0.75), or toward cultivation temperature (T_C_) (angle=8.8°, Rayleigh test= p<0.05, r=0.54; Fig. 3C). In a linear magnetic field of earth-strength (0.65 Gauss), animals showed a statistically significant migratory direction (angle=186.3°, Rayleigh test= p<0.01, r=0.68). In contrast, under no-field (angle=126.5°, Rayleigh test= p=0.77 r=0.16) and current controls (angle=179.9°, Rayleigh test: p=0.10, r=0.42) animals showed no statistically significant heading (Fig. 3C). These centroid-based trajectory results are consistent with the final animal position data used to calculate the orientation indexes above (Fig. 2).

**Figure 3.**
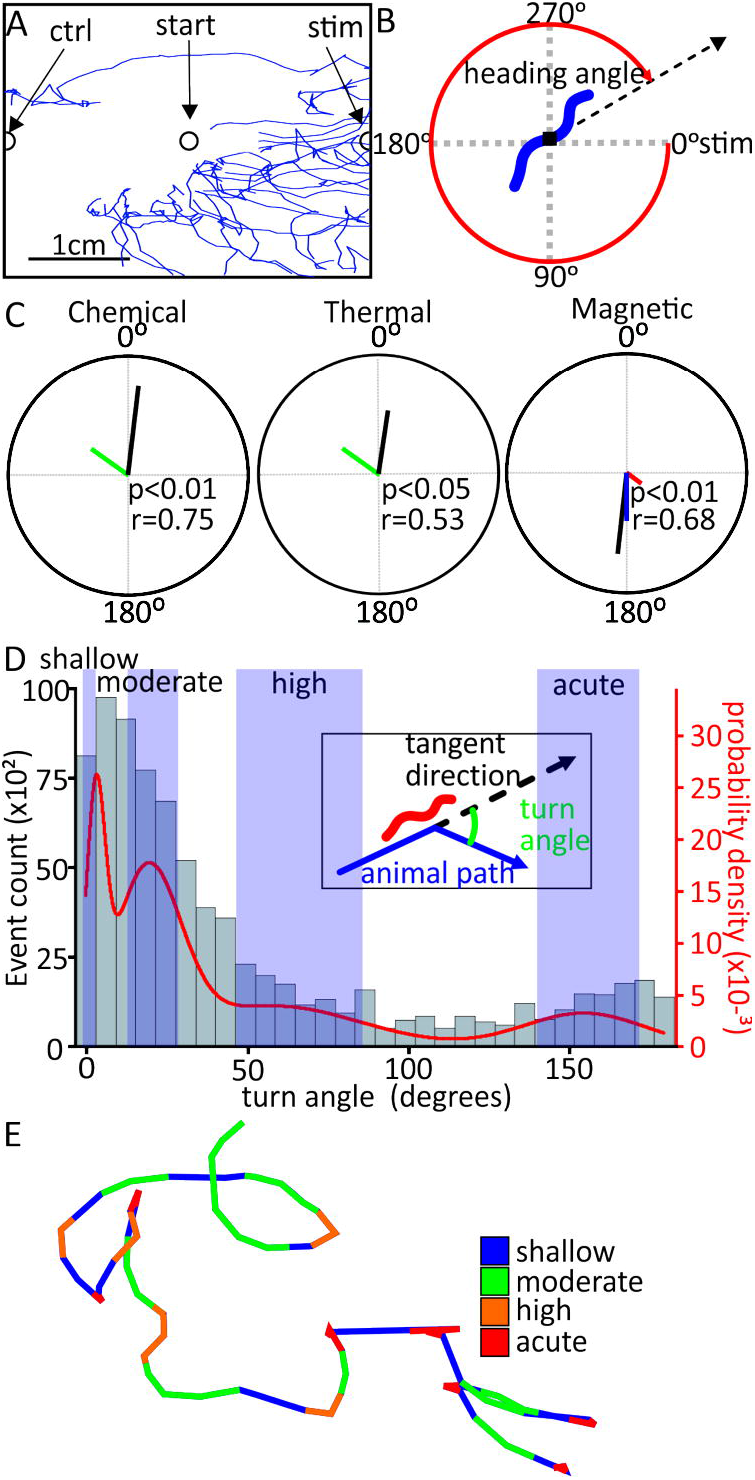
*C. elegans* perform different turning behaviors to maintain preferred migratory headings. **(A)** Sample of centroid tracks for animals migrating in a linear magnetic field. Animals began in center start position (open circle). **(B)** Heading angle (red arc) was defined as migration direction (dashed black arrow) relative to stimulus direction (defined as 0°). For magnetic orientation, 0° corresponded to the direction of magnetic north. **(C)** Circular plots of mean population headings (black lines). Radius vector length corresponded to fraction of the population moving along that heading. Control animals (green lines) did not show any significant headings. During chemotaxis (and thermotaxis), animals oriented their headings toward the attractant (or cultivation temperature, T_c_). In the presence of a linear magnetic field of earth strength (0.65 Gauss), worms displayed a significant migratory preference. Animals in a *no field* (red line), or *current control* (blue line) did not show significant headings. Significance was determined by Rayleigh’s test for non-uniformity. **(D)** Histogram showing the distribution of animal turning angles during navigation across all stimuli. Angles are measured from 0°-180°. Inset: Turning angle (green arc) was defined as change in angle of the animal path (blue arrow) and continued forward locomotion in the tangent direction (dashed black arrow). A Gaussian mixed model (solid red curve) shows the probability density estimates (right axis) for turn angles separated into four categories by clustering analysis. Blue rectangles show boundaries of each category used to define discrete behaviors; shallow angle turns: 0.0°-4.9°, moderate angle turns: 13.0°-26.9°, high angle turns: 46.0°-81.6°, and acute angle turns: 139.8°-170.7°. **(E)** Representative animal trajectory with turn angles highlighted with corresponding behaviors.

### Unbiased analysis of animal turn angles identifies behavioral components used during navigation

*C. elegans* locomotion can be decomposed into distinct ***behavioral components*** (Croll, 1975). These include straight forward motion (runs), steering toward the stimulus (weathervanes), single large amplitude turns (deep bends), turns where animal bend head to tail (omega bends), or backwards travel (reversals). Because the definition of these behaviors varies across studies and is based on animal postural changes (rather than the resulting trajectory changes), we focused our analysis on centroid trajectory changes. We used an unbiased approach to identify and define several behavioral components as extracted from the complete population of trajectory changes of all animals, in all assays, in our dataset. We thus fitted a mixed multiple gaussian model to the turn angle distribution extracted from automatically generated animal centroid coordinates (n=81,106 turns, N=66 assays). To measure the magnitude of turn angles, rather than handedness of the turns, angles were measured from 0°-180°. Behavioral components were defined using the interquartile range (IQR) of each peak from the gaussian fit (Fig. 3D). As described in the methods, turn angles were calculated as the angle between the animal’s trajectory defined across three successive time points (Fig. 3D, insert). Our analysis identified four distinct behavioral components: ***shallow angle turns***=0.0-4.9°, ***moderate angle turns***=13.0-26.9°, ***high angle turns***= 46.0-81.6°, and ***acute angle turns***=139.8-170.7° (Fig. 3E). These changes in trajectory have been previously associated with postural changes known as runs, weathervanes, deep bends, and reversals/omega bends respectively (Croll 1975, Iino and Yoshida, 2009, Vidal-Gadea et al., 2012). We elected to use a different nomenclature (introduced above) to reflect the fact that our definition is based on centroid trajectory changes, as opposed to body or postural changes. Our choice of measuring and reporting angles in a 0-180 scale (rather than 0-360 scale) assumes that worms are as likely to crawl on the left side of their bodies, as they are to crawl on the right side of their bodies. This assumption could potentially lead to the generation of artifactual turn distributions if animals in fact biased their crawling to one of their sides preferentially. We therefore double checked and confirmed the categories identified by our analysis by taking handedness of the turns into account. We observed equivalent turn angle distributions mirrored around 0° (data not shown).

### Validation of behavioral analysis using chemotaxis in C. elegans

Our initial velocity analysis indicated that strategies for thermo- and magnetotaxis were distinct from chemotaxis and control animals. Knowing that our turn angle analysis can separate at least four different categories, we set out to determine if animals use different turning strategies when orienting to different stimuli. These differences could manifest as changes in behavioral component rate, spatial distribution, or handedness (e.g. toward vs away stimulus). Because it is a thoroughly studied behavior, we used chemotaxis to test the acuity of our algorithm.

We compared the probability density distribution of turn angles in control and during chemotaxis but found no difference between chemotaxis and controls (Fig. 4A, top panel). Similarly, we found no difference in the mean rate of behavioral components carried out between chemotaxis and control (Fig. 4A, bottom panel). However, when we measured where animals were located during the assay using the probability density estimate (PDE) of animal positions, we found a significant bias towards the side of the attractant during chemotaxis (Fig 4B). No such trend was present in control, consistent with our earlier observation that animals migrated towards the attractant.

**Figure 4.**
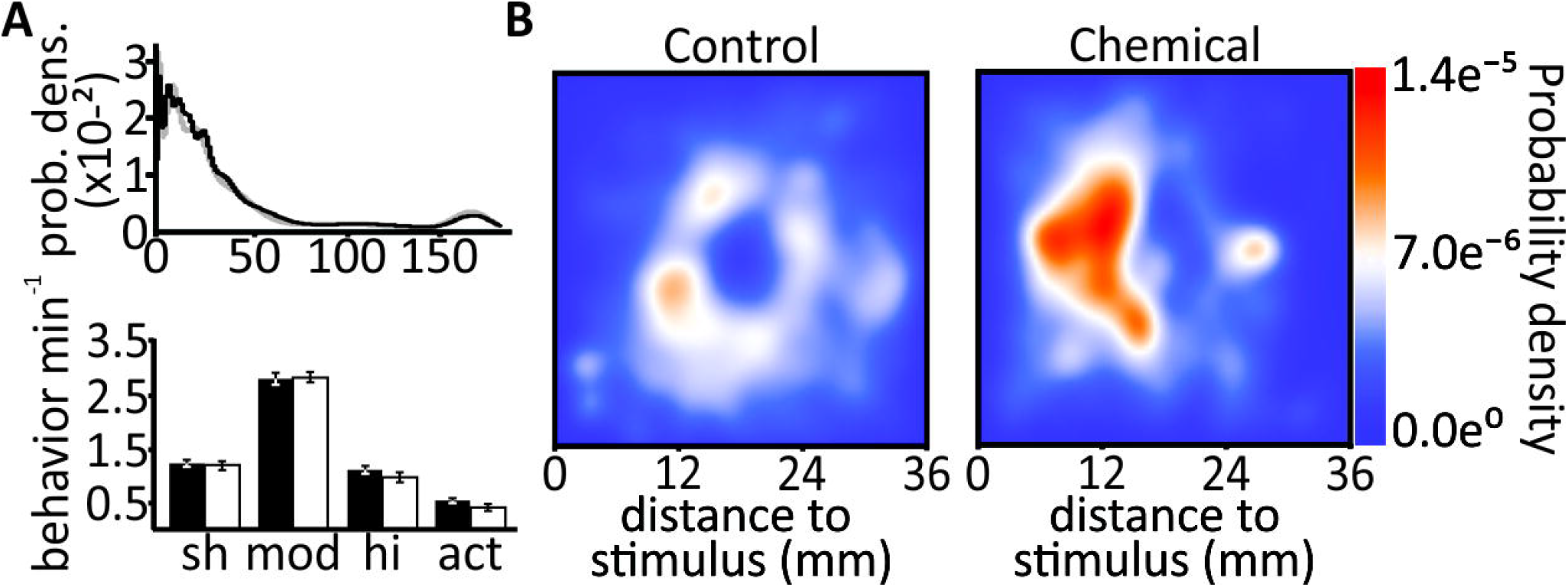
Orientation behaviors displayed under control and chemical stimulus conditions. **(A)** Top panel: the probability density estimate (PDE) of turn angle distributions for animals orienting in the absence of stimulus (gray trace), and in the presence of a chemical stimulus (black trace). Both stimulus conditions show nearly identical turn angle probability density distributions. Bottom panel: mean rate of behavioral turn events during control (black bars) and chemotaxis experiments (white bars) shows that animals did not alter their behavioral rates during chemical orientation. sh=shallow angle turn, mod=moderate angle turn, hi=high angle turn, act=acute angle turn. **(B)** Heat map PDE of animal positions (x-y coordinates) relative to their distance to the stimulus showing low (blue) and high (red) probability of animal spatial distribution. Distance from the stimulus (mm) is shown along the x-axis, with 0mm representing the stimulus. PDEs show probability distribution of animals migrating toward the stimulus during chemotaxis, whereas migrating in the absence of a stimulus are evenly distributed around the center start position.

Since *C. elegans* suppress reorientations to maintain course when travelling toward increasing attractant concentrations (Pierce-Shimomura et al., 1999; 2005), we predicted a reduction in reorientation probabilities (e.g. high and acute angle turns) as animals approached the attractant. While control animals performed acute angle turns and high angle turns uniformly over the assay space (Fig 5A), we observed a significant decrease in the number of acute angle turns as chemotaxing animals approached the stimulus (Fig 5B; Pearson’s coefficient: 0.27, N=10, p<0.01, Pearson product-moment).

**Figure 5.**
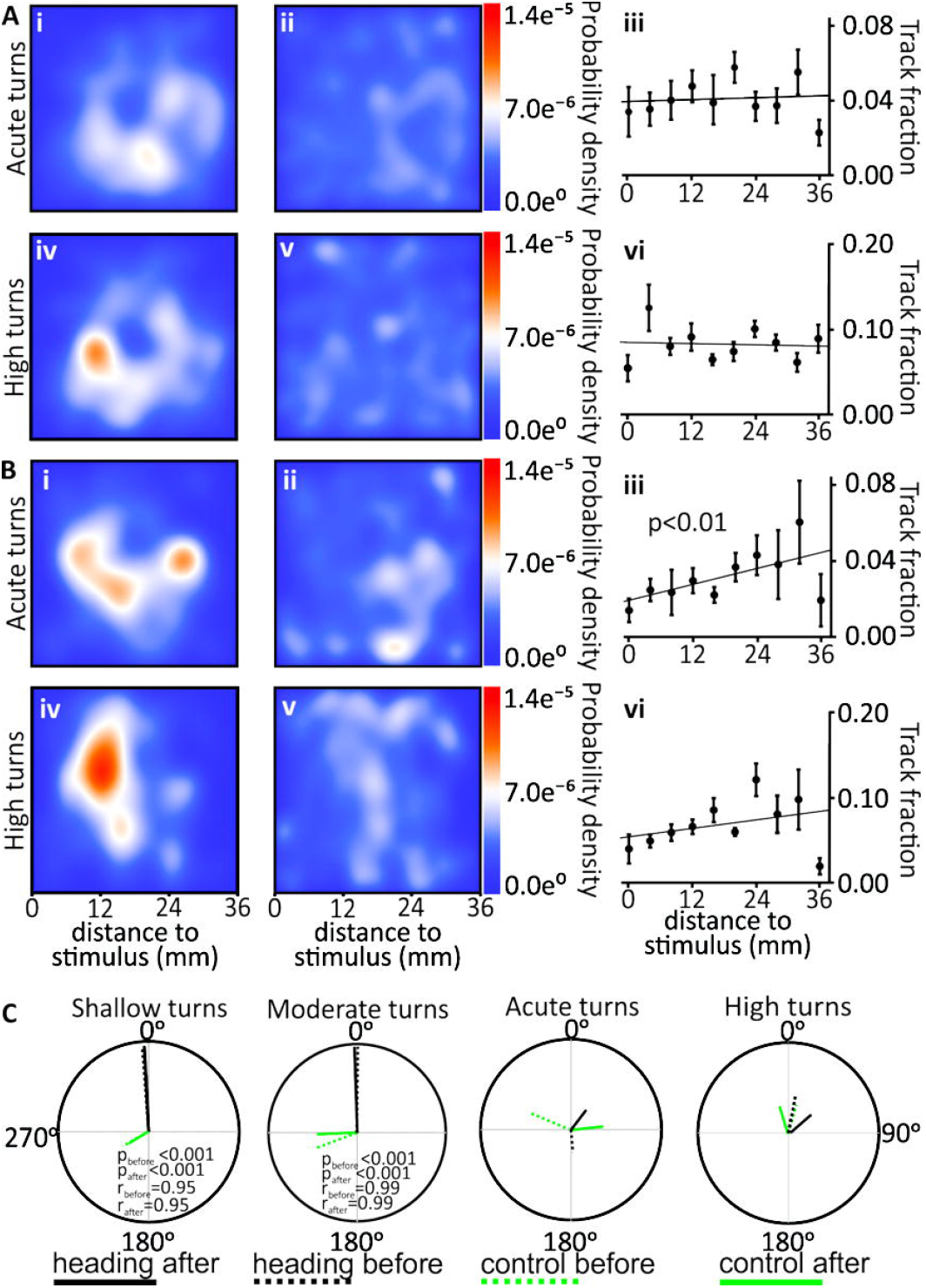
Validation of unbiased behavioral analysis through chemotaxis. **(A)** Probability density estimates (PDEs) for detected acute angle turns (i) for animals in non-stimulus controls. Normalizing of acute angle turns to their positional density revealed no spatial bias in the performance of these turns (ii), or their distribution along the x axis calculated as the mean fraction of track that were acute turns in each bin (iii, mean±sem). Similar results were obtained for high angle turns (iv-vi). **(B)** PDEs of animals performing acute angle turns (i-iii) or high angle turns (iv-vi) during chemotaxis. (i) PDE shows animals perform acute turns with higher probability during migration toward the stimulus. (ii) PDE of acute turns normalized to animal positional density reveals suppression of acute angle turn near the stimulus. (iii) Linear correlation showed acute turns significantly decreased near the stimulus. (iv-vi) PDE for high angle turns over chemotaxis assay space. (iv) Animals performed high turns during migration to the stimulus but, unlike acute angle turns, high angle turns were not suppressed near the stimulus (v). (vi) Track fraction analysis confirmed that animals did not alter high turn probability over the assay space. **(C)** Mean headings before and after different turning behaviors. 0° corresponds to stimulus directed locomotion. Radius vectors represent mean headings before (dashed lines), and after (solidlines) a behavior was performed. Behavioral headings showed animals performed shallow angle and moderate angle turns when their trajectory was significantly aligned with the stimulus. In contrast, animals performed acute angle, and high angle turns when their trajectories were random. Performance of these turns did not improve their heading significantly. Significance determined by Rayleigh’s test for non-uniformity.

Because *C. elegans* biases the direction of turns during chemotaxis towards the stimulus source (Pierce-Shimomura et al., 1999; 2005), we assessed if performance of behavioral components similarly improved an animal’s heading toward a stimulus. To do this, we compared animals heading before and after performing each behavioral component. We hypothesized that components contributing to align an animal with its stimulus would produce an improved, non-random, post-performance heading when compared to headings just prior to the behavior. We therefore compared post-performance headings to random distributions (that would be predicted if the behavior was independent of the stimulus).

As expected, control animals showed no significant alignments towards any direction for any behavioral components under consideration, either before or after their performance (Fig. 5C). Chemotaxing worms consistently moved in the direction of the stimulus. The performance of shallow angle turns did not significantly impact their heading (before shallow angle turns: angle=355.6°, Rayleigh test= p_before_<0.001, r_before_=0.95; after shallow angle turns: angle= 357.1°, Rayleigh test= p_after_<0.001, r_after_=0.95). The same was true for moderate angle turns (before: angle= 0.77°, Rayleigh test= p_before_<0.001, r_before_=0.99; after: angle= 358.05°, Rayleigh test= p_after_<0.001, r_after_=0.99; Fig. 7). Both moderate angle turns (Watson-Williams test; F(1,19)=0.49; p=0.49), and shallow angle turns (Watson-Williams test; F(1,19)=0.03; p= 0.87) did not significantly improve the heading of these animals. We did not observe a directional bias for high angle turns (angle=33.9°, Rayleigh test= p_before_=0.38, r_before_=0.31; angle= 88.3°, Rayleigh test= p_after_=0.60, r_after_=0.23) or acute angle turns (angle= 175.3°, Rayleigh test= p_before_=0.54, r_before_=0.25; angle= 37.2°, Rayleigh test= p_after_=0.47, r_after_=0.28, Fig. 5C). This was unexpected because a moderate turning bias towards chemical attractants has been previously described (Pierce-shimomura, 1999). Our analysis was based on mean assay headings, therefore our conflicting results could be due to an absence of turning bias, or alternatively, to reduced sensitivity in our analytical approach. We reanalyzed our data by treating headings before and after high angle turns, and acute angle turns, as individual events rather than using mean heading values. Because analysis of individual events were not based on mean headings, we report individual turns as a linear probability distribution measured from 0° to ±180° (Fig S2). This analysis produced a significant heading after acute angle turns (angle=-161.22°, Rayleigh test= p_before_ =0.23, r_before_=0.06; angle= 35.26°, Rayleigh test= p_after_<0.05, r_after_=0.087, Fig. S2). These results suggest that our behavioral detection algorithm is able to capture the majority of the behavioral strategies displayed during chemotaxis described by the literature.

### Worms thermotaxing up a gradient rely on directionally biasing and increasing the rate of high turns

We next investigated behavioral strategies for thermotaxis, when animals oriented to their cultivation temperature, T_c_. *C. elegans* uses two distinct methods for thermal orientation depending on its position relative to T_C_. When placed above their T_C_, animals suppress reorientations when moving along decreasing temperature to continue toward T_C_. Additionally, they bias the direction of turns toward cooler temperature (Ryu and Samuel, 2002; Clark et al., 2007; Zariwala et al., 2007). When placed below their T_C_, as in our assays, worms only bias the direction of turns toward increasing temperature, but do not suppress turning probability (Luo et al., 2014). We set out to determine if our analysis would be sensitive enough to detect the directional bias from high turns and acute turns during positive thermotaxis. The distribution of turning angles indicated a larger occurrence of high and acute turns than for controls (Fig. 6A). An analysis of the rate at which behavioral components occurred revealed overall significant changes from control animals (One-way ANOVA, N=43, F(3,39)=38.81, p<0.001). Specifically, we observed an increased acute turn rates (1.34±0.09 min^−1^, p<0.05, Holm-Sidak) over controls (0.55±0.05 min^−1^). Similarly, Kruskal-Wallis ANOVA on ranks (H(3)=18.73, p<0.001) showed a significant increase in high turn rates (1.59±0.07 min^−1^, p<0.05, Dunn’s post hoc) relative to controls (1.07±0.07 min^−1^). A One-way ANOVA on shallow turns (N=43, F(3,39)=8.97, p<0.001) and moderate turns (N=43, F(3,39)=14.96, p<0.001) showed a reduction in the rates of both shallow (0.86±0.07 min^−1^, p<0.05, Holm-Sidak) and moderate turns (1.83±0.10 min^−1^, p<0.05, Holm-Sidak) (Fig. 6B). Hence, *C. elegans* oriented to increasing temperatures by increasing the rate of acute and high turns and by reducing those of shallow and moderate turns.

**Figure 6.**
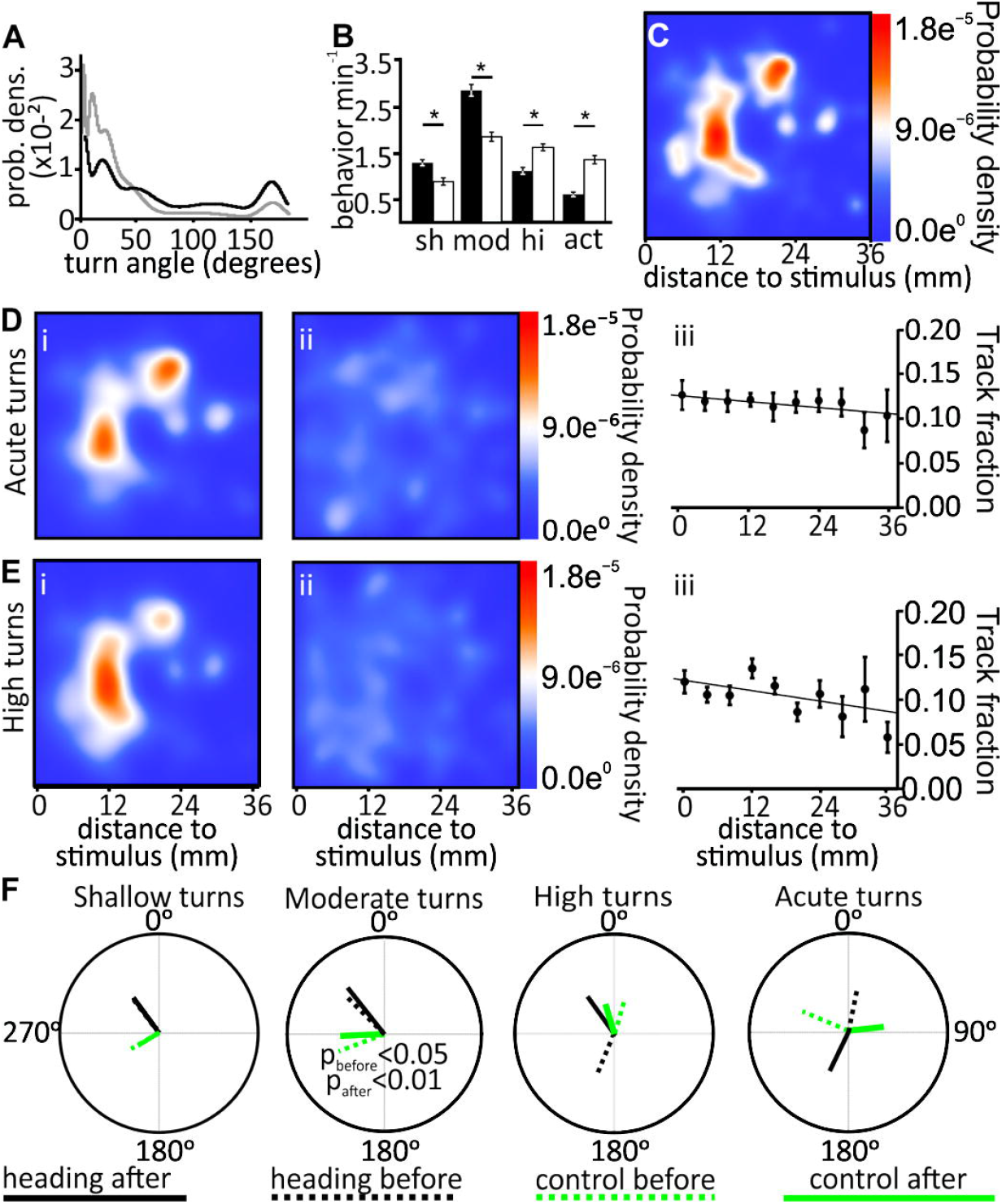
Orientation strategies for thermotaxing animals. **(A)** Probability density estimates of turn angles for thermotaxing animals (black trace) compared to control animal distribution (gray trace). **(B)** Animal behavioral rate under control conditions (black bars) and during thermotaxis (white bars). Thermotaxing *C. elegans* (white bars) increased the rate of high angle turns and acute angle turns compared to animals orienting in the absence of a stimulus (black bars). Conversely thermotaxing animals reduced the rate of shallow turns and moderate turns (asterisks correspond to p<0.05). **(C)** PDE heat map of animal positions during thermotaxis. Animal positions were distributed with the highest probability toward their T_C_. **(D)** Spatial distribution of all detected acute turns during thermotaxis. (i) PDE of acute angle turns during migration toward their T_C_. (ii) Normalized (to positional density) PDE of acute angle turns. (iii) Linear correlation of acute turn angle track fraction with distance to stimulus. **(E)** Spatial distribution of high angle turns shows similar trend as acute angle turns. (i) Worms performed high turns during migration to T_C_, however their normalization to their spatial distribution showed no spatial bias in their performance (ii), which was confirmed by their linear correlation (iii). **(F)** Mean headings of thermotaxing animals before and after performance of each type of turn. Animals traveling significantly aligned with their T_C_ performed moderate angle turns. However, animals did not show a significant stimulus alignment either before or after performing shallow angle, high angle, or acute angle turns. Significance determined by Rayleigh’s test for non-uniformity.

Worms showed an increased probability density near their T_C_, indicating directional migration toward their T_C_ (Fig. 6C). Unlike chemotaxis, we did observe a change in the probability of acute angle turns over the assay space for thermotaxing animals (Fig. 6D). Similarly, the probability of performing high angle turns also remained uniform independent of the distance to the stimulus (Fig. 6E). These results are consistent with previous reports that *C. elegans* does not suppress reorientations during positive thermotaxis.

We next quantified the contribution to migration toward a thermal stimulus by measuring headings before and after the performance of individual behavioral components. Moderate angle turns improved the heading of migrating animals (angle=314.9°, Rayleigh test = p_before_<0.05, r_before_ =0.54; angle= 321.3°, Rayleigh test =p_after_<0.01, r_after_= 0.6). However, neither shallow angle turns (angle= 322.84°, Rayleigh test = pbef=0.10, r_before_=0. 42; angle= 323.75°, Rayleigh test = p_after_=0.08, r_after_= 0.44) (Fig. 6F), high angle turns (angle= 324.5°, Rayleigh test= p_after_=0.08, r_after_=0.44), nor acute angle turns (angle= 205.8°, Rayleigh test= p_after_=0.09, r_after_=0.43) improved trajectories. As with chemotaxis, the conservative nature of our analysis prevented us from detecting some of the subtler behavioral changes described by the literature to take place during thermotaxis. Reanalysis of the data pooling across assays similarly detected a significant contribution for high angle turns (angle=-20.3°, Rayleigh test= p_after_<0.05, r_after_=0.04; Fig. S2B).

### *C. elegans* magnetic orientation does not rely on turn-suppression strategies

Unlike chemotaxis or thermotaxis, orientation strategies underlying magnetotaxis in *C. elegans* are underscribed. We turned to identify behavioral components underlying magnetic orientation. We started by determining if the probability of specific turns changed in comparison to control. Indeed, the turn angle distribution during magnetotaxis was different from control animals, and also markedly distinct from those of chemo- and thermotaxis (Fig. 7A). This suggested that animals were using a unique orientation strategy to orient to the magnetic field. We wondered if changes in behavioral rate might account for the unique angle distribution observed for magnetotaxis. We found that, as in chemotaxis but in contrast with thermotaxis, magnetotaxing animals displayed no behavioral rate change compared to controls (Fig. 7B). Since magnetotaxis appeared to not rely on changing the rate of behavioral components, we investigated if it relied on suppressing turning probability relative to the magnetic field. Such strategies are useful when comparing changes in stimulus intensity and is employed by chemotaxing animals as shown above. However, because the earth’s magnetic field has uniform intensity, a strategy relying on turn suppression relative to changing stimulus intensity seemed unlikely. We first analyzed where in the arena animals carried out the individual behavioral components. Our previous analysis revealed that animals migrated opposite the magnetic field direction (Fig. 3C). Consistent with this observation, we found animals congregated away from the side towards which the magnetic vector pointed (i.e. southwardly, Fig. 7C). We next investigated whether animals altered turn production when moving along their preferred direction. We found that worms performed acute angle turns with equal probability across the assay area (Fig. 7C). Pearson’s product moment analysis confirmed that there was no significant correlation between acute (Pearson’s coefficient=0.01, p=0.94, pearson’s product moment) or high angle turn performance (Pearson’s coefficient= 0.10, p=0.36, pearson’s product moment) and animal spatial location on the assay (Fig. 7D). Therefore, magnetotaxing worms appear to not rely on neither the frequency, nor the spatial distribution of turning events in order to orient to magnetic stimuli.

**Figure 7.**
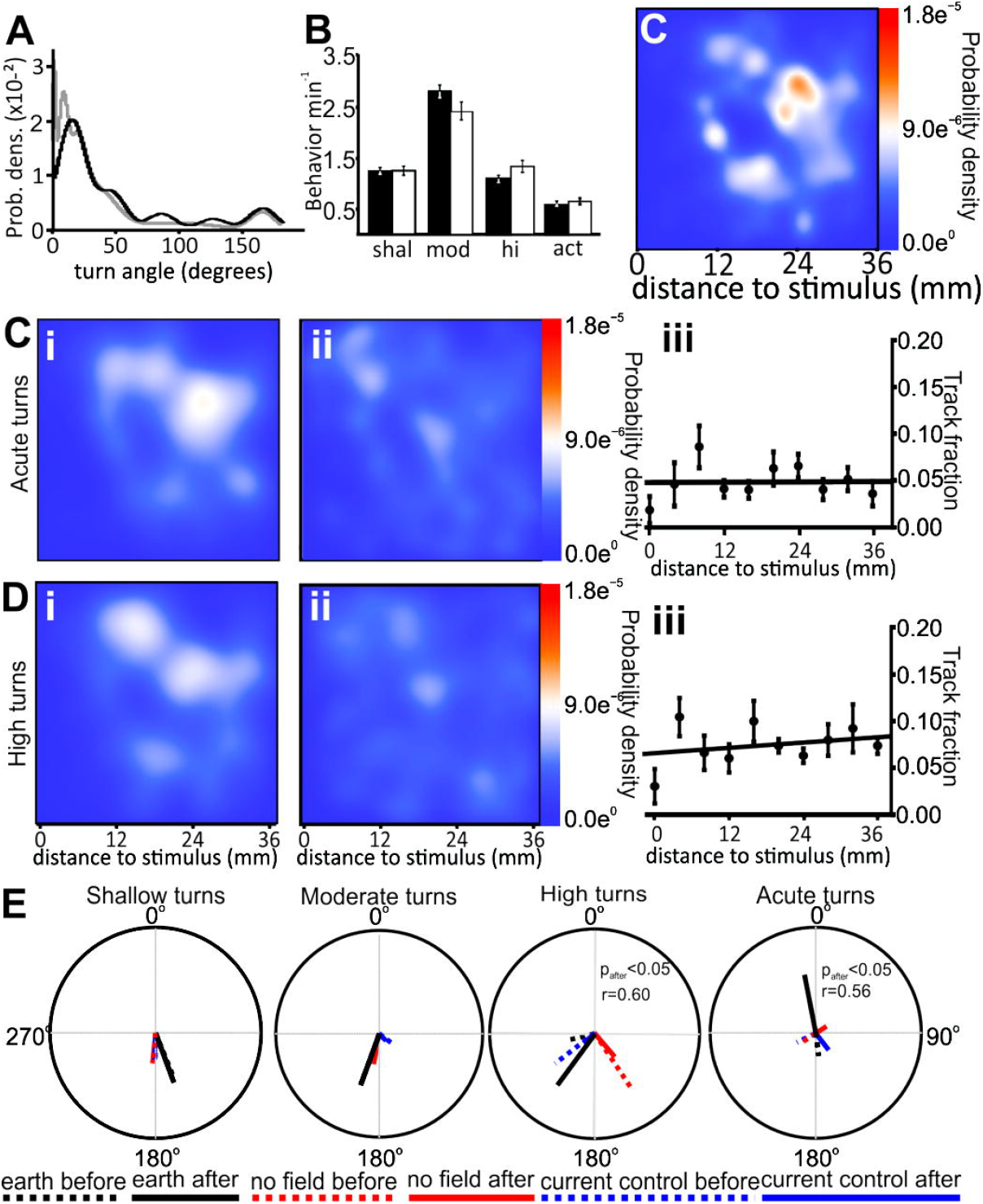
Orientation strategies for magnetotaxing animals. **(A)** Probability density estimates turn angles during magnetotaxis (black trace) and control animals (gray trace). **(B)** The behavioral rate during magnetotaxis (white bars) and control experiments (black bars) did not vary significantly. **(C)** Density distribution of animals during magnetotaxis shows them migrating in the opposite direction of the field. **(C)** Probability density of acute angle turns over the assay (i). (ii) Normalization of acute turns showed animals performed acute angle turns with equal probability relative to distance to stimulus. (iii) Linear correlation of acute angle turn track fraction with distance to stimulus confirmed that animals perform acute turns uniformly over the assay during magnetotaxis. **(D)** (i) Probability density estimates of high angle turns. Animals performed high angle turns as they migrated. (ii) high angle turn normalization to animal density shows equal probability of high turning over the assay, confirmed by linear correlation of track fractions (iii). **(E)** Headings of magnetotaxing animals before and after performance of each turn behavior. Animals showed significant headings in the direction of migration following high angle turns, indicating these turns facilitated magnetic orientation (solid black line). Animals displayed significant headings in the direction of the magnetic field vector after performing acute turns (solid black line). Significance determined by Rayleigh test for non-uniformity of circular data.

### *C. elegans* orientation to magnetic fields involves turn angle modulation

We analyzed if the performance of the different turning behaviors components identified by our analysis improved orientation towards the preferred migratory direction. We found robust turning biases for acute angle and high angle turns (Fig. 7E). While the initiation of an acute angle turn was not associated with animals traveling in any particular direction (angle=172.6°, Rayleigh test= p_before_=0.62,r_before_=0.22), following the execution of an acute angle turn animals exited traveling at an angle consistent with the direction of the imposed magnetic field (angle= 349.1°, Rayleigh test = p_after_<0.05, r_after_=0.56). Additionally, we found that animals performing high angle turns also displayed a statistically significant heading after performing this turning maneuver (angle= 256.2°, Rayleigh test= p_before_=0.55 r_before_=0.25; angle= 216.3°, Rayleigh test = p_after_<0.05, r_after_=0.60). However, unlike acute angle turns, performance of high angle turns aligned animals at an angle to the magnetic field consistent with their preferred direction of migration. In the absence of the imposed magnetic stimulus (no field, and current controls) performance of acute angle turns (angle= 157.9°, Rayleigh test= p_before_=0.11, r_before_=0.47; angle= 159.8°, Rayleigh test =p_after_= 0.08, r_after_=0.50) or high angle turns (angle = 200.1°, Rayleigh test= p_before_=0.06, r_before_=0.53; angle = 200.2°, Rayleigh test = p_after_=0.05, r_after_= 0.53) did not significantly impact migratory direction. Thus, orientation to magnetic stimuli by *C. elegans* appears to involve turn angle modulation during the performance of acute angle, and high angle turns.

## Discussion

The ability of *C. elegans* to detect and orient to a diverse range of stimuli makes it a compelling system to investigate how animals alter search strategies to environmental stimuli. Our unbiased approach to categorize turning behaviors allowed us to compare behavioral strategies across different sensory conditions. In contrast to chemical and temperature stimuli, magnetic fields are vectorial and have uniform intensity and direction at organismal scales. Thus, magnetic fields do not provide changes in intensity like chemical or thermal gradients. We found that magnetic orientation in *C. elegans* relies on a strategy in which animals modulate high amplitude turns to align their trajectory with an imposed magnetic field, and modulate the amplitude of high angle turns to align their trajectories with their preferred migratory direction. These results indicate that *C. elegans* employs distinct orientation strategies when migrating to chemical, thermal, or magnetic stimuli.

### Use of chemotaxis and thermotaxis to validate unbiased behavioral analysis

This study aimed to characterize the orientation strategy *C. elegans* uses to orient to magnetic fields, and to determine whether behavioral strategies employed for orientation to this vectorial stimulus differs from that used in other sensory modalities. The orientation strategy used to locomote towards chemical stimuli is arguably the best characterized, with much of the neural circuitry, and molecular players involved in chemical detection and orientation mapped (Gray et al., 2005; Dunn et al., 2004; Larsch et al., 2015; Luo et al., 2014). A chief strategy during chemotaxis is turn suppression, which effectively biases the direction of turns toward increasing concentration to avoid moving down concentration gradients (Pierce-Shimomura et al., 1999; 2005). Our approach was able to detect the majority of the strategies associated with chemotaxis and thermotaxis in *C. elegans* (Luo et al., 2014).

Unbiased detection algorithms have been used before to detect locomotor defects of mutant phenotypes and describe discrete changes between animal states (Yemini et al., 2013). While these approaches made great contributions to identifying locomotor deficits, they generally lack the ability to determine orientation strategies as they do not identify corresponding behavioral components. We designed an unbiased approach to identify and quantify behavioral components thus minimizing the potential for subjective experimenter bias. Our approach has the ability to directly identify and characterize behavioral components under different sensory conditions. It allowed us to classify discrete turn angle ranges to define behavioral components as the animal locomotes. This method could be applied to identify differences in orientation strategies that extract information from other vectorial and graded stimuli, such as polarized light, or local odor trails (El Jundi et al., 2016, Wehner et al. 2016). Our approach is, however, not without caveats. Our algorithm used animal centroids to obtain unbiased and robust measurement of trajectory changes at the expense of finer (postural) resolution. We therefore consider the present a first approach to measuring and comparing diverse behaviors which should then be followed by a more in depth analysis of identified strategies.

### *C. elegans* uses distinct behavioral strategies to orient in magnetic fields

The main goal of this study was to study how *C. elegans* alters its behavior during orientation to vectorial stimuli in the absence of spatial or temporal gradients. In contrast to graded stimuli, such as chemicals or temperature, the earth’s magnetic field is vectorial and uniform, meaning that it is linear and lacks intensity or directional changes at organismal scales. Consequently, rather than relying on changes in intensity, many magnetosensitive species orient relative to vectorial properties of the magnetic field, such as its polarity or inclination. For example, loggerhead turtles orient making use of local geomagnetic field conditions (Lohmann et al., 2001; Fuxjager et al.,2011, Putman et al., 2015). However, how these animals acquire relevant magnetic field information, or which behavioral strategies they perform to navigate in the field remains to be worked out.

Like other magnetosensitive species, *C. elegans* migrates at a preferred direction to the magnetic field that is state dependent, with fed and hungry animals electing to migrate at opposite angles (Vidal-Gadea et al., 2015). Here we provide the first insights into how worms, and perhaps other species, might engage behavioral strategies in order to sample the magnetic field, and determine adaptive trajectories based on their internal states. Interestingly, we found that worms performing acure amplitude turns preferentially aligned with the direction of the imposed field (Fig. 7E). This behavior could be consistent with a sampling (or resetting) of the magnetic field prior to assuming the preferred migratory direction. Indeed, recent work on migratory white throated sparrows showed that these birds use polarized sunlight at dusk and down to recalibrate their internal compass prior to initiating their migratory flight (Muheim et al., 2009). Similarly, the migratory Bogong moth uses the magnetic field to calibrate its headings relative to visual cues by periodically sampling the magnetic field (Dreyer et al., 2018). Periodic body alignment with the geomagnetic field vector is thought to underlie compass calibrations and likely serve to orient animals relative to less reliable stimuli (Begall et al., 2013; Bianco et al. 2019).

In addition to using acute angle turns to align their trajectories with an imposed magnetic field, *C. elegans* uses high angle turns to align their paths with the preferred migratory trajectory (Fig. 7E). Together, these two distinct behavioral strategies might allow worms to discern the direction of the earth’s magnetic field, and then select the appropriate migratory direction depending on the organism’s internal state.

*C. elegans* detects both thermal and magnetic information using the same pair of AFD neurons. Despite detection through the same neuron, our data show a divergence in the behavioral strategies used for these two different modalities. While usually thought of as a feature of distributed neuronal networks (Follmann et al., 2018), there is already precedence for the divergence of behavior selection by AFD neurons with regards to orientation to increasing or decreasing temperature (Clark et al., 2006; Kimura et al., 2004;Luo et al. 2014; Ramot et al.,2008a, b). In thermotaxis this divergence occurs within the same sensory modality, and avoids potential processing conflicts by virtue of not occurring at the same time (animals cannot be in higher and lower than cultivation temperature at once). In the case of magnetic orientation, the behavioral strategies for sampling orienting to the earth magnetic field, and to orient to temperature gradients are not in conflict because they involve different behaviors. Practically, this could mean that animals could orient and migrate to both temperature gradients and the magnetic field of the earth simultaneously as long as the two stimuli were aligned.

In our previous work we suggested that worms might use the magnetic field of the earth to migrate vertically in their environment (Vidal-Gadea, 2015). Temperature gradients in the soil provide reliable vertical information, but are also ambiguous as the direction of the gradient cycles through the day and across the seasons (hotter up during the day/summer, hotter down at night/winter). In contrast, the magnetic field of the earth provides a reliable cue which perhaps coupled with temperature sensation would allow the AFD neurons to unambiguously guide animals as they navigate in their environment. Combining magnetic field detection with other modalities is in fact a common proposed mechanism for its function as exemplified by the proposed detectors in the retina of birds (Mouritsen and Hore, 2012; Günther et al., 2018.).

Much remains to be learned about magnetic orientation in *C. elegans*. How animals detect and transduce this information, or how it is then processed by downstream effectors is not yet known. The behavioral, cellular, and molecular tractability of *C. elegans* make this a useful organism to decipher how animals interact with the earth’s magnetic field. The findings in this and other studies suggest that the lessons we may learn from studying this tiny worm will likely transfer to many other magnetosensitive species.

## Acknowledgements

Some strains were provided by the Caenorhabditis Genetic Center, which is funded by NIH Office of Research Infrastructure Programs (P40 OD010440) and the National Bioresource Project of Japan. Funding was provided by ISU URG program to AGVG and by NSF grant to AGVG: 1818140. We would like to thank Dr. Bill Perry for extensive help with R.

---

**Figure S1.** Animals orienting to magnetic stimuli show directional bias but do not alter the rate of any turning behaviors compared to magnetic controls. **(A)** Worms migrating to a magnetic field showed a directional bias (N=14, 0.45±0.08) compared to animals migrating to no field (N=14, 0.14±0.02) or under current control conditions (N=17, 0.13±0.03). Asterisk indicates significance p<0.05). **(B-E)** Turning behaviors for animals orienting to magnetic fields. There were no significant differences between *no field* or *current control* conditions for any turning behaviors. Significance was determined using ANOVA on ranks using Dunn’s correction.

**Figure S2**. Turning bias before and after acute angle turn and high angle turn performance. Headings in these histograms are pooled for all turning behavior events across respective assays. Headings are measured from 0° to ±180° with 0° corresponding to motion toward stimulus and ±180° corresponding to headings away from stimulus through oppositely biased (left or right) turns. **(A)** Analysis of headings for animals orienting to a chemical attractant. Acute angle turn PDE (top row) and high angle turn PDE (bottom row) before (left column) and after (right column) each behavior. Headings for animals moving down the concentration gradient were improved toward the stimulus following acute turns. **(B)** Heading analysis during thermotaxis of acute angle turn probability (top row) and high angle turn probability (bottom row), before (left column) and after (right column) each respective behavior. High angle turns improved headings of animals toward T_C_. Significance determined by Rayleigh Test.

